# Calcium nanodomains in spindles

**DOI:** 10.1101/404715

**Authors:** Guolong Mo, Ruizhen Li, Zachary Swider, Yong Tao, Katsuhiko Mikoshiba, William M. Bement, X. Johné Liu

**Affiliations:** Ottawa Hospital Research Institute, Ottawa Hospital General Campus, 501 Smyth Road, Ottawa, K1H 8L6. Canada; Department of Biochemistry, Microbiology and Immunology. University of Ottawa; Department of Zoology, University of Wisconsin-Madison, 1117 West Johnson Street, Madison, WI 53706, USA; Laboratory for Developmental Neurobiology, Brain Science Institute, RIKEN, Saitama 351-0198, Japan; Department of Obstetrics and Gynaecology, University of Ottawa

**Author notes:** GM and RL contributed equally to this work. **Correspondence:** Johné Liu Ottawa Hospital Research Institute 501 Smyth Road Ottawa, K1H 8L6 Canada; Bill Bement Department of Zoology University of Wisconsin-Madison Madison, WI 53706, USA.

## Abstract

The role of calcium signaling in specific events of animal cell meiosis or mitosis (M-phase) is a subject of enduring controversy. Early efforts suggested that increases in intracellular free calcium ([Ca^2+^]_i_) promote spindle disassembly ^1, 2^ while subsequent work suggested that global [Ca^2+^]_i_ increases trigger nuclear envelope breakdown, spindle assembly, the metaphase-anaphase transition, and cytokinesis ^3-6^. However, further studies led to the conclusion that elevation of [Ca^2+^]_i_ either has no role in these events, plays a permissive role in these events, or functions as an auxiliary signaling pathway that supplements other mechanisms ^7^. One potential explanation of the controversy is that specific M-phase events might depend on highly localized increases in [Ca^2+^]_i_, variously referred to as microdomains ^8^ or nanodomains ^9^, as proposed recently ^10^. Such domains are hypothesized to arise from rapid shuttling of calcium between closely positioned sources and sinks, rendering them potentially difficult to detect with traditional dyes and largely insensitive to slow chelators such as EGTA ^9^. Here a novel microtubule-binding calcium sensor—TubeCamp--was used to test the hypothesis ^10^ that spindles are associated with calcium nanodomains. TubeCamp imaging revealed that spindles in *Xenopus* eggs, *Xenopus* embryos, and HeLa cells were all associated with calcium nanodomains at the spindle poles. Calcium nanodomains also formed in spindles assembled in cell extracts and at the center of monopolar spindles, suggesting that they are a basic feature of spindle self-assembly. Disruption of calcium nanodomains via perturbation of inositol-1,4,5-trisphosphate signaling or rapid chelation of [Ca^2+^]i resulted in spindle disassembly in vivo and vitro. The results demonstrate the existence of spindle-associated calcium nanodomains and indicate that such domains are an essential and common feature of spindles in vertebrates.

---

To overcome the limitations of soluble calcium reporters, we developed a genetically encoded probe designed to detect microtubule-proximal increases in [Ca^2+^]_i_. This probe, dubbed TubeCamp, comprises the calcium-sensitive derivative of GFP, GCamp3 ^11^, fused with the microtubule-binding domain of ensconsin (EMTB) ^12^. GCamp fluorescence emission increases upon calcium binding ^11^ while fusions of EMTB with fluorescent proteins have been used in mammalian ^13^, amphibian ^14, 15^ and invertebrate ^14^ cells to label microtubules. To determine whether TubeCamp reports on microtubule-proximal elevated [Ca^2+^]_i_, TubeCamp and R-Geco, a calcium-sensitive reporter protein ^16^ were expressed in *Xenopus* oocytes which were then wounded to elicit a local [Ca^2+^]_i_ increase and microtubule reorganization ^17^. Before wounding, the oocyte cortex displayed low levels of both R-Geco and TubeCamp fluorescence (Fig 1A); immediately after wounding, both R-Geco and TubeCamp fluorescence increased sharply in a circular region around the wound (Fig. 1A). While the global pattern of TubeCamp fluorescence paralleled that of R-Geco in space and time (Fig. 1A, A’), the TubeCamp signal was distinctly filamentous. That the filamentous structures detected by TubeCamp were microtubules was determined by wounding experiments using TubeCamp in combination with 2X-mCh-EMTB (mCh-EMTB; Fig. 1B): mCh-EMTB labeled all cortical microtubules before and after wounding; TubeCamp fluorescence was sharply elevated after wounding only on microtubules within 10 µm of the wound (Fig. 1B, B; and C). TubeCamp also reported on microtubule-proximal increases in [Ca^2+^]_i_ in somatic cells in developing embryos, as shown by wounding one epithelial cell, which triggers and increase in [Ca^2+^]_i_ in neighboring epithelial cells (Supplemental Fig. 1; ^18^).

**Figure 1.**
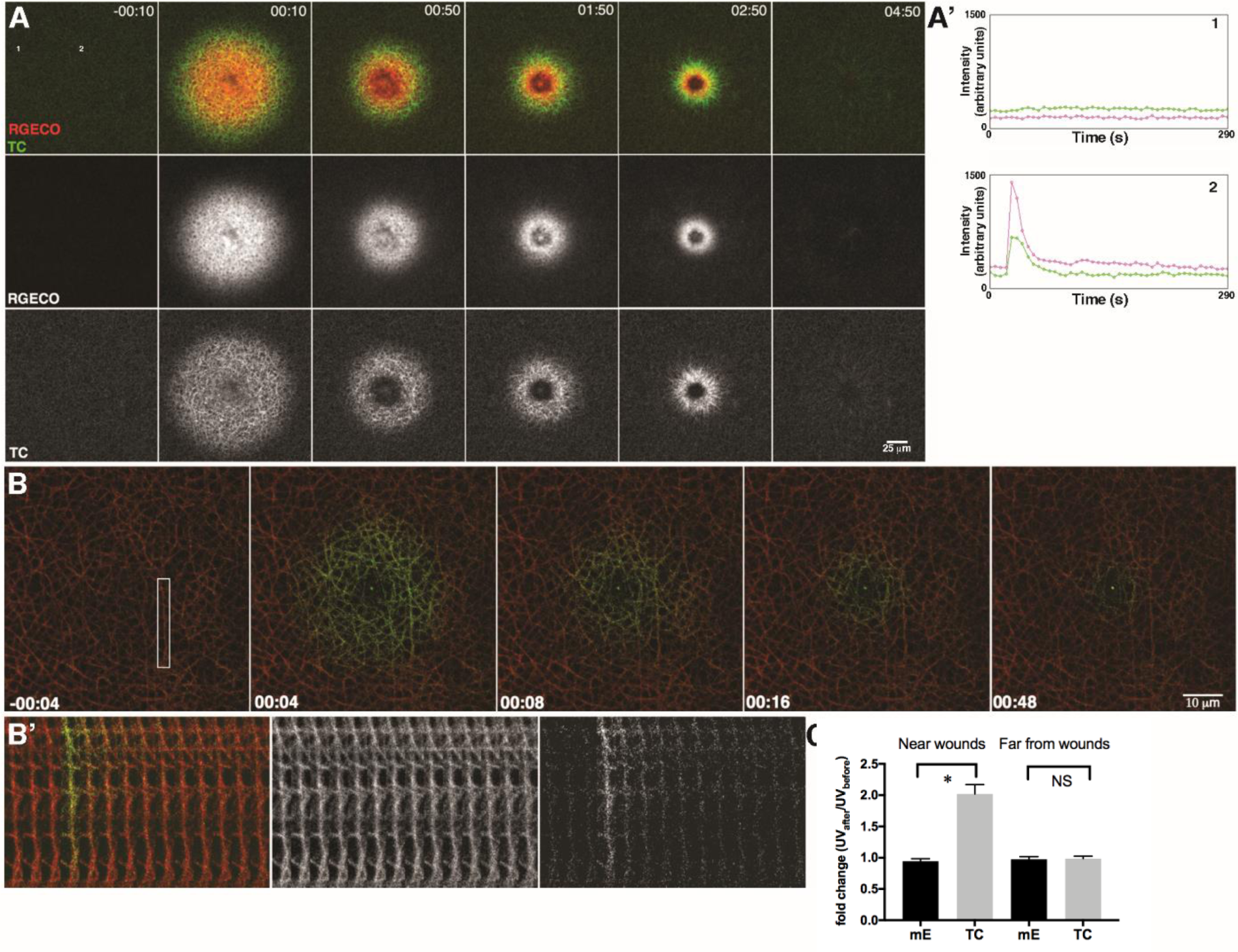
TubeCamp specifically reports on microtubule-proximal [Ca^2+^]_i_. **A.** Still frames from confocal movie of wounded *Xenopus* oocyte showing total intracellular free calcium (revealed by RGECO; red) and microtubule-proximal intracellular free calcium (revealed by TubeCamp [TubeCamp]; green). Time in min:sec; sample wounded at 0 sec. **A’**. Fluorescence intensity plots of regions indicated as 1 and 2 in first panel in A with 1 (top plot) distal to the wound and 2 (bottom plot) proximal to the wound. The temporal patterns of elevated [Ca^2+^]_i_ (RGECO; red) and TC (green) closely parallel each other. **B.** Still frames from confocal movie of wounded *Xenopus* oocyte showing total microtubules (revealed by mCh-EMTB; red) and microtubule-proximal [Ca^2+^]_i_ (TubeCamp; green). Time in min:sec; sample wounded at 0 sec. Prior to wounding (−00:04) cortical microtubules are labeled with mCh-EMTB but not TubeCamp. Shortly after wounding (00:04) microtubules within ∼ 20µm of the wound acquire green fluorescence; this fluorscence disappears as wound heals. **B’**. Montage showing enlargement of area boxed in first frame of B at 4s intervals. Wounding occurs between 3^rd^ and 4^th^ panel and is accompanied by local, microtubule-associated increase in green (TubeCamp) but not red (mCh-EMTB) fluorescence. **C.** Quantification of relative mCh-EMTB and TubeCamp fluorescence on microtubules before and after wounding and within 10 µm or farther than 30 µm from wound. Microtubules within 10 µm of wound undergo a significant increase in TubeCamp but not mCh-EMTB fluorescence. Results are mean +/- SD; * indicates p < 0.0001; n=11.

To determine whether *Xenopus* egg meiotic spindles are dependent on calcium nanodomains ^10^, TubeCamp was co-expressed with mCh-EMTB (Fig. 2A) or rhodamine-tubulin (not shown). While cortical TubeCamp signal remained low for most of meiosis, in prometaphase TubeCamp fluorescence began to rise at the spindle pole closest to the cortex (Fig. 2A). As the spindle became bipolar, TubeCamp fluorescence was evident at both poles and much more concentrated there than mCh-EMTB (Fig. 2A). In addition, interpolar TubeCamp fluorescence was observed that, again, clearly differed from that of total microtubules. Interpolar TubeCamp fluorescence disappeared prior to metaphase, while the polar signal was maintained until after anaphase (Fig. 2A) when it disappeared (not shown) until meiosis II during which it reappeared at the poles (Supp. Fig. 2). That TubeCamp responds to calcium in the oocytes was demonstrated by the dramatic increase of TubeCamp signal at metaphase II spindle (Supp. Fig. 2) within minutes of inducing the fertilization-specific cortical calcium wave ^10, 19^.

**Figure 2.**
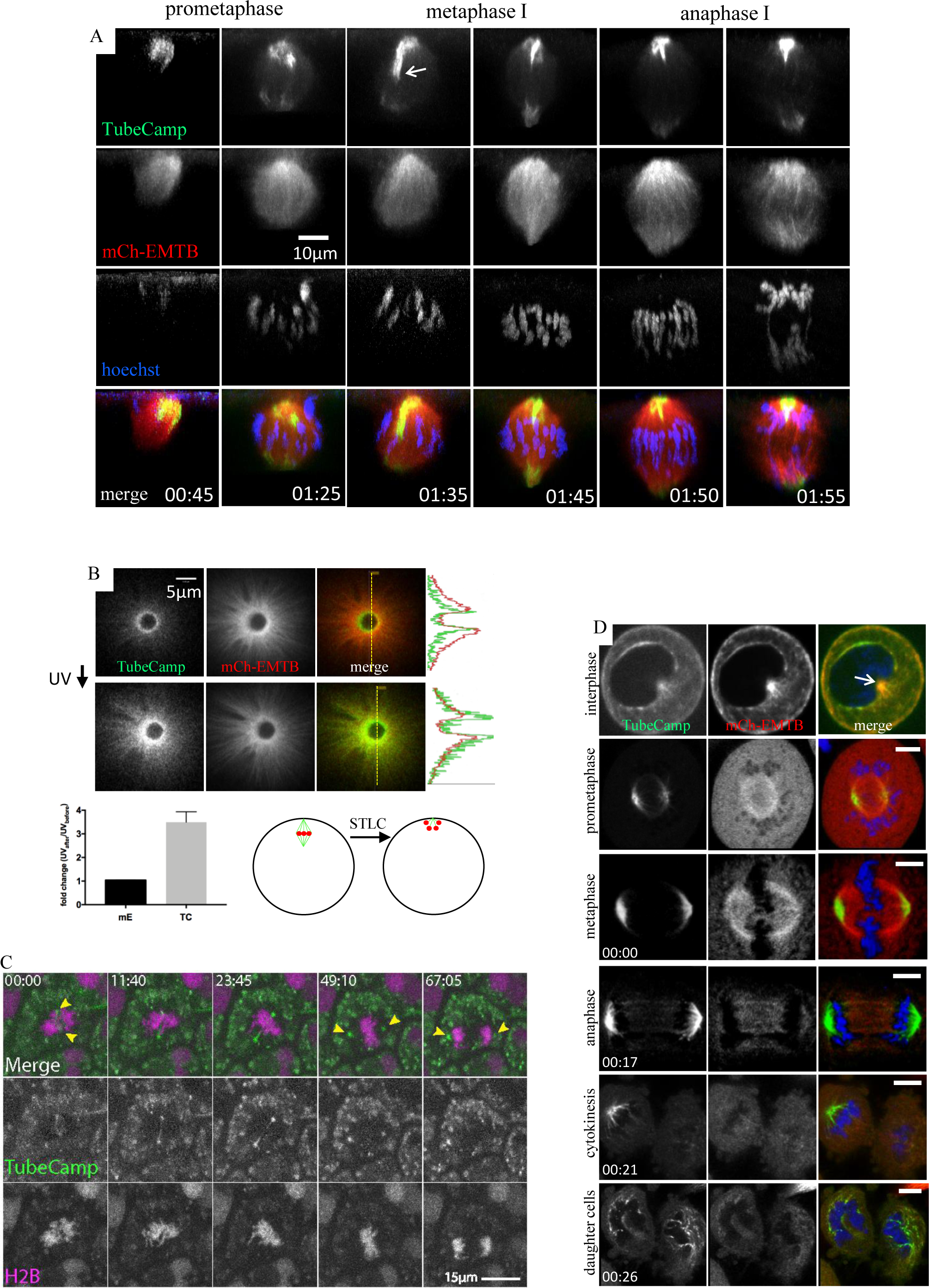
TubeCamp signal in meiotic and mitotic cells. **A.** 3D rendering of confocal z stacks of *Xenopus* oocytes expressing TubeCamp and mCh-EMTB at the indicated stage of oocyte maturation. Time in hr:min after GVBD, TubeCamp signal is focused at spindle poles and along distinct bundle of interpolar microtubules (arrow). All except the left column (00:45) are from the same oocyte. **B.** Live confocal images of monopolar spindle before (top row) and immediately after (bottom row) UV uncaging of IP_3_; corresponding fluorescence intensity line scans on right. TubeCamp (TC) fluorescence, but not mCh-EMTB (mE) fluorescence, significantly increased upon uncaging (graph; p<0.0001; n=9). Schematic depicts the locations of a bipolar spindle (those shown in A) or a monopolar spindle (here) in intact oocytes (green: microtubules; red: chromosomes). **C.** Live confocal of *Xenopus* neurula epithelial cells coexpressing TubeCamp (green) and mCherry-H2B (magenta). TubeCamp signal is concentrated along the spindle axis and as discrete foci at the poles (yellow arrows). Non-spindle associated signal (seen in both channels) is the result of yolk autofluorescence. Time is in min:sec (Movie 2). **D.** Confocal images of HeLa cells in interphase (arrow: centrosome), and the various stages of mitosis. Images of the four bottom rows are acquired from the same cell. Time (hr:min) is from the beginning of imaging of this cell. Spindle is tilted in cytokinesis image, thus only one pole is visible. TubeCamp signal is concentrated at the spindle poles during mitosis and near decondensing chromosomes shortly after cytokinesis. Scale bars: 5µm.

To further substantiate the presence of spindle-based Ca^2+^ transients, we imaged oocytes using a mobile Ca^2+^ indicator, Oregon green 488 BAPTA-2 (OG-2). Consistent with the results obtained with TubeCamp, OG-2 signal significantly increased at the spindle assembly site in time course similar to that shown by TubeCamp (Supp. Fig. 3; Movie 1). In contrast to the results obtained with TubeCamp, OG-2 revealed only diffuse-spindle-associated signal, likely due to its relative mobility.

**Figure 3.**
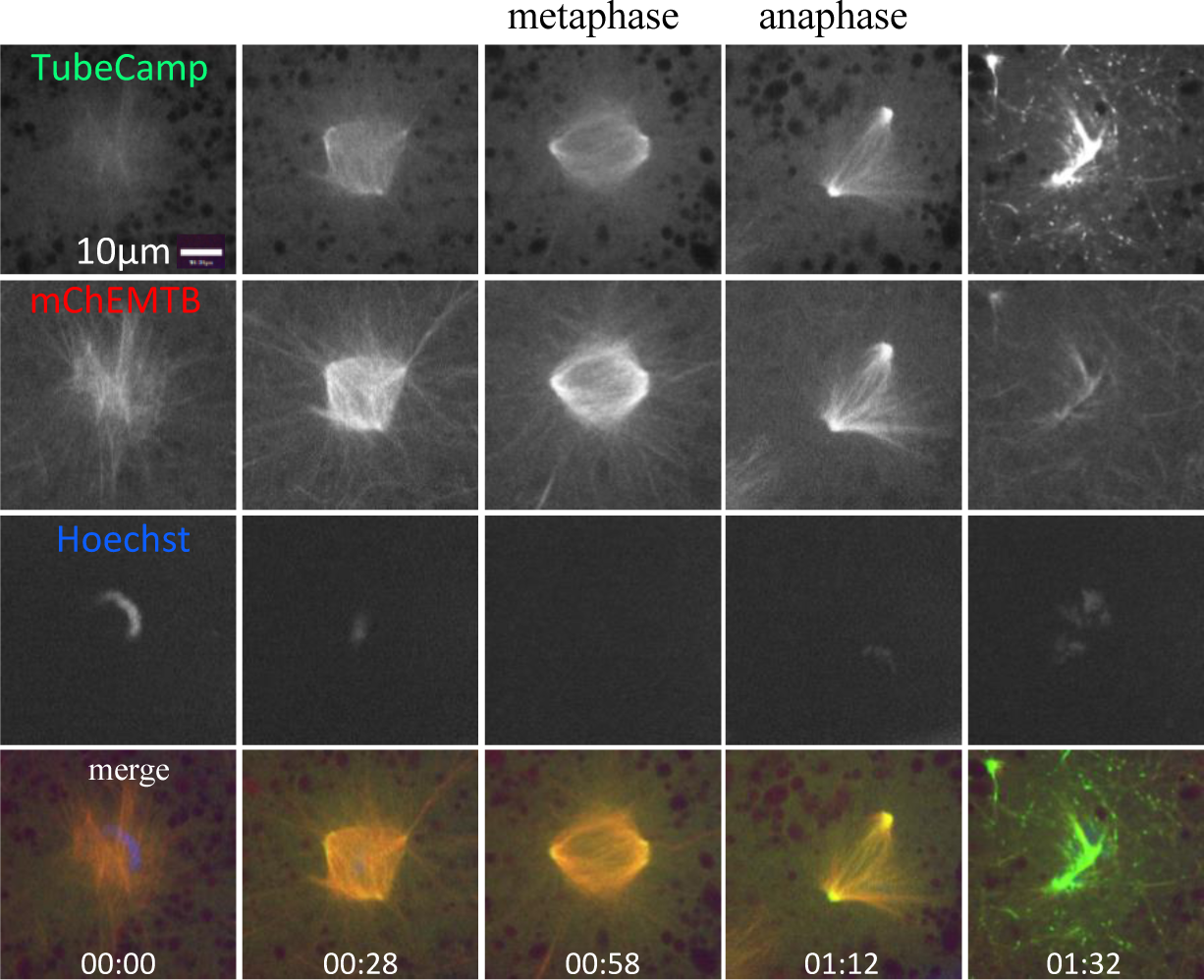
TubeCamp signal during spindle assembly in cell extracts. Confocal time series of a bipolar spindle in extract of GVBD oocyte expressing TubeCamp and mCh-EMTB. Time (hr:min) is from the beginning of imaging. TubeCamp signal is concentrated at spindle poles. Individual TubeCamp nanodomains became visible after anaphase (Movie 3).

The spindle pole localization of TubeCamp signal was also evident in oocytes induced to form monopolar spindles in the presence of the kinesin 5 inhibitor S-trityl L-cysteine (STLC) ^20^. In these oocytes, TubeCamp signal concentrated at the ring-shaped monopole (Fig. 2B, top row). To further confirm the specificity of the TubeCamp probe, oocytes with monopolar spindles were subject to uncaging of IP_3_ ^21^. This manipulation resulted in an immediate and dramatic increase of TubeCamp signal but not that of mCh-EMTB signal (Fig. 2B).

To determine whether calcium nanodomains associate with mitotic spindles, *Xenopus* gastrula expressing TubeCamp and mCherry-Histone were analyzed. As in meiotic spindles, TubeCamp signal was elevated at each of the spindle poles and along a microtubule or bundle of microtubules running from pole to pole (Fig. 2C; Movie 2).

To extend this discovery to mammalian mitosis, we expressed TubeCamp with mCh-EMTB (Fig. 2D) or mCh-α-tubulin (Supp. Fig. 4) in HeLa cells. As in *Xenopus* meiotic and mitotic spindles, TubeCamp signal was observed at the two spindle poles from prometaphase to cytokinesis (Fig. 2D). Following cytokinesis, TubeCamp signal persisted around the chromosomes where individual TubeCamp foci were observed in the two daughter cells (Fig. 2D).

**Figure 4.**
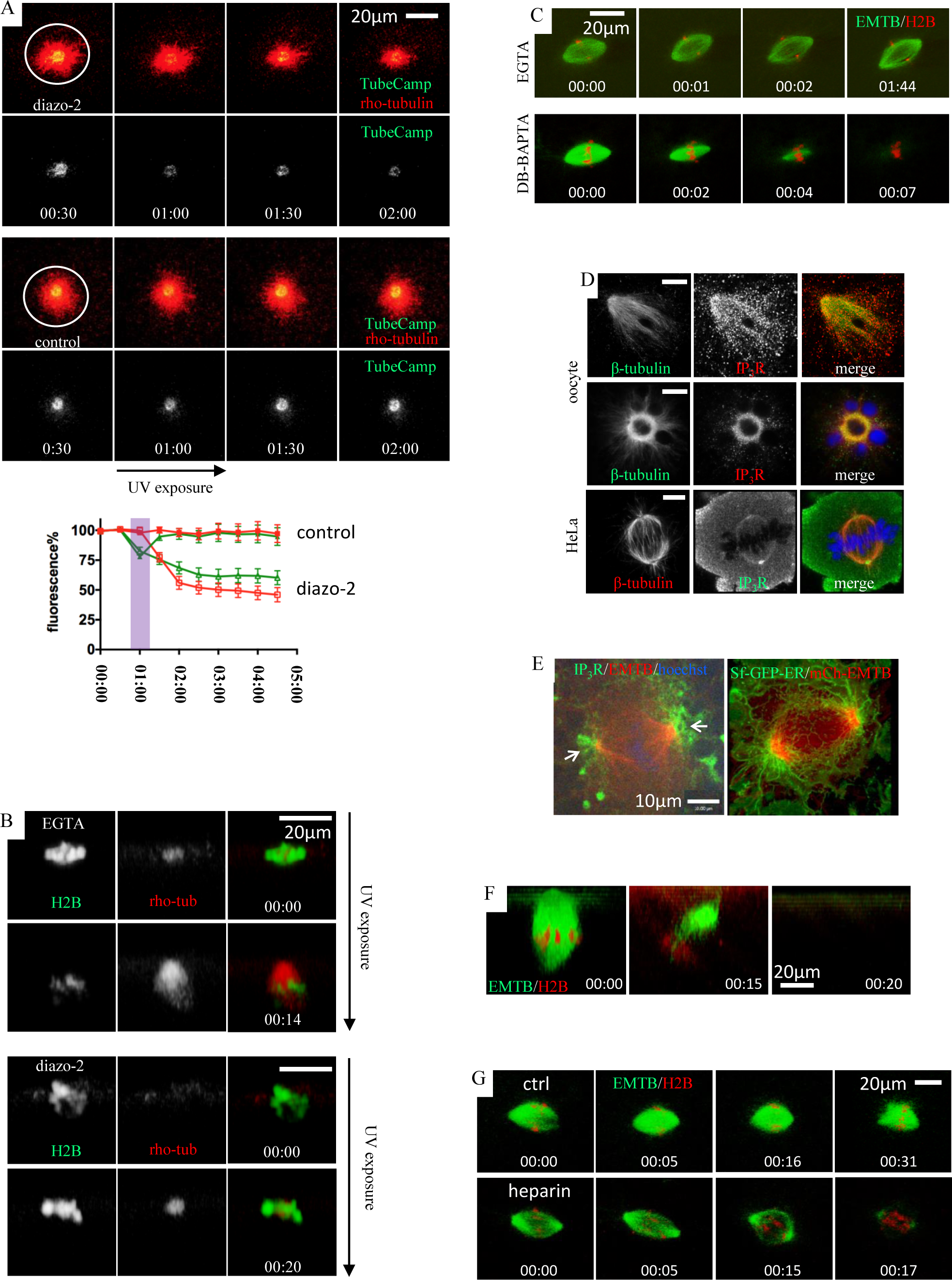
Calcium nanodomains are required for spindle formation and stability. A. Confocal images of monopolar spindles subjected to uncaging of diazo-2 (caged BAPTA). Top: following uncaging by UV exposure, both TubeCamp signal and microtubules disappear; Bottom, in control samples exposed to UV but not injected with diazo-2, TubeCamp signal and microtubules persist (Movie 4). The graph summarizes relative fluorescence (means±SEM; green and red represent TubeCamp and rho-tubulin respectively) at the indicated time points from 9 oocytes in each of control and diazo-2 groups. Purple strip depicts UV exposure which slightly photobleached TubeCamp. B. Confocal images of normal meiotic spindles allowed to recover from microtubule depolymerization. DNA labeled with GFP-Histone H2B (H2B) and microtubules with rhodamine tubulin. Top: following inactivation of colcemid in the presence of EGTA, the spindle reforms. Bottom: following inactivation of colcemid in the presence of BAPTA (simultaneously photolyzed from diazo-2), the spindle does not reform (Movie 5). C. Confocal images of metaphase spindles formed in extracts treated with EGTA (2.5 mM) or dibromo-BAPTA (2.5 mM). DNA labeled with mCh-H2B; microtubules labeled with GFP-EMTB. Dibromo-BAPTA, but not EGTA, causes spindle dissolution. D. Immunofluorescence analysis of IP_3_ receptor distribution in normal meiotic spindles (top row), monopolar spindles (middle row) or HeLa cells (bottom row). Receptor is particularly concentrated at spindle poles. E. Left: Distribution of eGFP-IP_3_ receptor (green), microtubules (mCh-EMTB; red) and DNA (Hoechst, blue) in extract-assembled spindle. Arrows: spindle poles. Right: Distribution of ER (Sf-GFP-ER; green) and microtubules (mCh-EMTB; red) in extract assembled spindle. F. Meiotic spindle in oocyte before (00:00) or 15 (00:15) or 20 (00:20) min after microinjection with heparin (200μg/mL). Microtubules labeled with GFP-EMTB (green); DNA with mCh-H2B (red). Heparin results in dissolution of the spindle. Time in hr:min. G. Control and heparin-treated (200µg/mL) extract spindles; microtubules labeled with GFP-EMTB (green); DNA labeled with mCh-H2B (red). Heparin causes disolution of the spindle. Time in hr:min.

Spindle calcium nanodomains might arise as a specific consequence of the normal three dimensional organization of the cell or they might develop as a basic feature of spindle self-organization ^22^. To distinguish between these possibilities, a micro-aspiration approach was devised that allowed spindle assembly in cell free extracts obtained from single *Xenopus* oocytes (Supp. Fig. 5). To examine Ca^2+^ nanodomains in this system, we added demembranted sperm to cytoplasm from GVBD oocytes expressing TubeCamp and mCh-EMTB. No distinct TubeCamp signal was observed at the early stage when an aster was seen (Fig. 3, 00:00). However, TubeCamp signal concentrated at the spindle poles as the spindle became bipolar (Fig. 3, Movie 3). Thus, calcium nanodomains are a basic feature of spindle self-organization.

To test the significance of the spindle nanodomains, oocytes treated with STLC (to induce monopolar spindles) and expressing TubeCamp and rhodamine tubulin were subjected to UV-uncaging of diazo-2, a caged BAPTA analogue ^10, 23^. Uncaging of diazo-2 resulted in dissipation of the spindle nanodomains and, in parallel, loss of spindle microtubules (Fig 4A; Movie 4).

As a complementary approach, meiotic spindle reformation after colcemid treatment was examined in the presence of EGTA or diazo-2. Spindles were first disassembled by colcemid treatment and the colcemid was then inactivated by UV exposure ^15^. In the presence of EGTA, the slow chelator, meiotic spindles reformed (Fig. 4B, EGTA, 00:00-00:14; Movie 5) but in the presence of diazo-2 they did not (Fig. 4B, diazo-2, 00:00-00:20; Movie 5), because UV photolysis also released the fast chelator BAPTA ^10^ in addition to inactivating colcemid ^15^ in the oocytes. As a further test of nanodomains, spindles were assembled using the microaspiration approach and then treated with either EGTA or dibromo-BAPTA ^10^. While spindles exposed to EGTA persisted for more than an hour, those exposed to BAPTA disassembled within minutes (Fig. 4C), as we have shown in intact oocytes ^10^.

To determine whether calcium stores are associated with spindles, immunofluorescence was used to monitor the location of the inositol-1,4,5-trisphosphate (IP_3_) receptor. Strikingly, the IP_3_ receptor was concentrated at the poles of normal meiotic spindles, monoastral spindles and HeLa cell spindles (Fig. 4D). Further, both the IP_3_ receptor and endoplasmic reticulum (ER) were concentrated at the poles of spindles assembled in extracts (Fig. 4E). As a functional test of these potential calcium stores, heparin, an IP_3_ receptor antagonist (5) was employed. Heparin caused rapid loss of spindle microtubules both in intact oocytes (Fig. 4F) and in extracts (4G).

In summary, the results provide the first direct visualization of calcium nanodomains in M-phase cells, show that such nanodomains are essential for spindle assembly and maintenance, and indicate that they are likely dependent on IP_3_-gated calcium stores. The results also show that in addition to forming at poles, the nanodomains are associated with a population of microtubules running from pole-to-pole that, to the best of our knowledge, has not been described before. More generally, the strategy employed here may be broadly useful for identifying other potential M-phase calcium micro or nanodomains via fusion of GCamp (or related calcium reporters) with nuclear envelope proteins, histone, or other proteins localized to sites of hypothesized calcium increase.

## Acknowledgement

We thank Dr. Chloë van Oostende-Triplet and Ms. Skye McBride of University of Ottawa Image Core for advice and assistance during this study. We thank Drs. Yixian Zheng and Rebecca Heald for advice on *Xenopus* egg extract preparation, and Dr. Wayne SR Chen for discussion. This work was supported by a research grant from Canadian Institute of Health Research (MOP 89973) to XJL and an NIH grant (GM52932) to WMB. GM is a recipient of a scholarship from China Scholarship Council, and a Designation 2020 Faculty of Medicine Scholarship of University of Ottawa.

## Methods

Rabbit polyclonal antibodies against IP_3_ receptor-1(H-80) are from Santa Cruz. Mouse monoclonal antibodies against IP_3_ receptors, 4C11, have been described ^24^. H-80 was combined with monoclonal anti-tubulin β (DM1B, ICN) ^25^ for oocyte immune-staining. 4C11 was combined with rabbit anti-tubulin β (Santa Cruz) for HeLa cell immune-staining. We used the MaxChelator program (https://web.stanford.edu/∼cpatton/maxc.html) to calculate the ratio of Ca^2+^ buffers over CaCl_2_ (EGTA:CaCl_2_=4:1; dibromo-BAPTA:CaCl_2_=10:1) to give the desired free Ca^2+^ concentration,∼140nM ^19^, in the calcium buffers used in extract spindle experiments.

### Construction of plasmid for TubeCamp

First, the entire GCaMP3 coding sequence (Addgene #22692) was excised using Bgl II and Not I. The fragment was treated with Klenow before being inserted into the Stu I site of pCS2+ vector ^26^, resulting the expression plasmid pCS2-GCaMP3 ^10^. To generate TubeCamp, we PCR-amplified the sequence coding for the microtubule-binding domain of E-MAP115, EMTB ^14^, using the following two primers (5’ and 3’ respectively): 5’-TATGAATTCACCATGGCAGTGCGAAGCGAAACA and 5’-TATGAATTCGAAGAGCCCTCAGGTGG. The amplified DNA was digested with EcoRI followed by being inserted into the EcoRI site of pCS2+GCaMP3, described above ^10^. The resulting plasmid, TubeCamp, expresses EMTB at the N-terminus followed by the original GCaMP3 coding sequence including its N-terminal poly-His tag ^11^. These cloning manipulations also created a seven amino acids insert (NSRDLAT) between EMTB and the initiating methionine of GCaMP3. The plasmid was linearized with Not 1 and transcribed in vitro using SP6 polymerase (Ambion kit).

### Oocyte isolation and injection

Oocytes were manually defolliculated ^27^ and were kept at 18 ^o^C in OCM medium (oocyte culture medium: 60% of L-15 medium (Sigma), supplemented with 1.07 g BSA per liter, mixed with 40% autoclaved water to yield the appropriate isotonic solution for amphibian oocytes). Manually defolliculated oocytes were used for all live cell imaging and cytoplasmic droplet experiments without further treatment. When oocytes were used for immunofluorescence experiments (Fig. 3A), manually deffoliculated oocytes were treated briefly with collagenase ^27^ to remove residual follicle cells ^28^ before use.

Manually defolliculated oocytes were injected with mRNA encoding various probes, as described in our previous publications ^29-31^. For plasmid DNA, we usually injected 1-3 nL of highly purified plasmid DNA dissolved in water (∼1mg/mL) into the germinal vesicle, which is located directly under the animal pole, measuring about 0.5mm in diameter ^27^. The injected oocytes were incubated in OCM for at least 6 hours (mRNA), or up to 2-3 days, before the addition of progesterone (1µM) to induce oocyte maturation.

### Live imaging and image analyses

Oocytes were monitored for GVBD (germinal vesicle breakdown, indicated by the appearance of a white maturation spot) every 10 min. GVBD oocytes were individually transferred to fresh OCM without progesterone and further incubated, until the time of fluorescence imaging or cytoplasm aspiration for in vitro spindle assembly (see later).

Oocytes were imaged in poly-lysine-coated glass bottom microwell dishes (MatTek Corporation, P35G-1.5-10-C) with a 60x oil objective on a Zeiss Axiovert with a BioRad 1024 laser scanning confocal imaging system ^30^, or a Quorum Spinning Disk confocal system. Time lapse image series were collected at various time intervals. Each time point volume was comprised of 15-30 image planes 1-3 µm apart. Image series were 3D-rendered using Volocity (version 6.3). Fluorescence quantification and co-localization analyses were performed using Volocity program. All time-series are made from images acquired from the same cell, unless otherwise indicated.

For super-resolution (LSM880 with AiryScan, Zeiss) imaging, cells were similarly imaged using a 60 x oil objective. Images were typically acquired in super-resolution x and y dimensions (40 x 40 nm) but much bigger z steps (0.5-1 μm). Super-resolution confocal z-stack acquisition in intact oocytes took an average of 5 min for each time point. Therefore it is not practical to acquire a complete series during oocyte maturation or mitotic cell cycle, due to the significant fluorescence photo-bleaching and possible photo-induced cell cycle disruption. Snapshot images or short time series during critical transition were typically employed. Images were processed using the ZEN 2.3 Lite program provided by Carl Zeiss.

For UV photolysis (on the MRC 1024 system), the oocyte animal pole was exposed to UV excitation (Chroma’s 11000V3, 350/50 nm; 100W mercury bulb) through the same 60x oil objective, and simultaneously subjected to confocal imaging ^15^. UV exposure time was controlled by an electronic shutter (LAMBDA SC, Sutter Instrument) using manufacturer’s program.

### Single cell extract spindle assays

All operations were performed in the glass bottom microwell dishes and covered with mineral oil. Demembranated sperm nuclei (500 nuclei/µL) ^32, 33^ were placed at the glass bottom of the dish in droplets ∼5 nL. In some experiments, Hoechst dye was added to sperm to a final concentration of 1 µg per mL before application to the glass. Oocytes, typically at the time of GVBD or MII, were placed under oil in the same dish with their animal pole facing up. A glass pipette attached to the microinejctor and with a tip-opening of ∼30 µm was then forced into the oocyte from the animal hemisphere. Negative pressure (“fill” function) was applied to slowly aspirate oocyte cytoplasm into the glass pipette (up to 300nL) followed by expelling the cytoplasm onto the sperm droplets using the “inject” function. Multiple cytoplasmic droplets, 50-100 nL, could be produced from one oocyte. The dish was then placed on the microscope for confocal imaging. When chemical inhibitors were used, they were delivered, via an on-stage microinjector, on top of the cytoplasmic droplet, furthest away from the spindle being imaged (which was at the bottom in our inverse microscope system). The inhibitors were delivered in volume less than 1/10 of the cytoplasmic droplet.

### HeLa cell methods

HeLa cells were transfected via Lipofectromine (ThermoFisher) according to manufacturer’s instruction. Transfection was carried out on poly-lysine-coated glass bottom dishes using the following components (per 3cm dish): 0.25μg of mCh-EMTB and 0.13µg of TubeCamp, 4μL of Lipofectamine, 2mL of OptiMEM medium plus 5% fetal bovine serum (FBS). The cells (∼90% confluent) were incubated in the transfection mixture for 6 hours followed by change into fresh α-MEM medium plus 5% FBS. Transfected cells were imaged either directly in the transfection dishes the next day, or were split into new glass bottom dishes and imaged in subsequent days. Prior to imaging, Hoechst dye was added to 1µg/mL to the cells for 5 minutes before changes into dye-free medium.

### Imaging Xenopus embryos expressing TubeCamp

Albino *Xenopus* eggs were obtained, fertilized, and de-jellied as described previously ^18^. Fertilized embryos were injected at the one cell stage with 16nl of a mixture containing 4ng/µl of mCherry-H2B mRNA and 20ng/µl of TubeCamp mRNA. Embryos were cultured for 3 days at 14°C before imaging. *Xenopus* embryo images were acquired on an Opterra Swept Field Confocal (Bruker) equipped with a 60x 1.4NA oil objective using a 60µm pinhole array and an Evolve Delta EM-CCD camera (photometrics). Because images were acquired using a multispectral filter set, bleed-through from the mCherry signal into the TubeCamp channel was corrected by subtracting 25% of each mCherry frame from each corresponding TubeCamp frame using FIJI (ImageJ). Both channels were registered for drift using the StackReg plugin in FIJI, and both channels were corrected for bleaching using the simple ratio adjustment in FIJI.

### Statistics

Data were analyzed using Student’s t-test (two tailed).

**Supplemental Figure 1.**
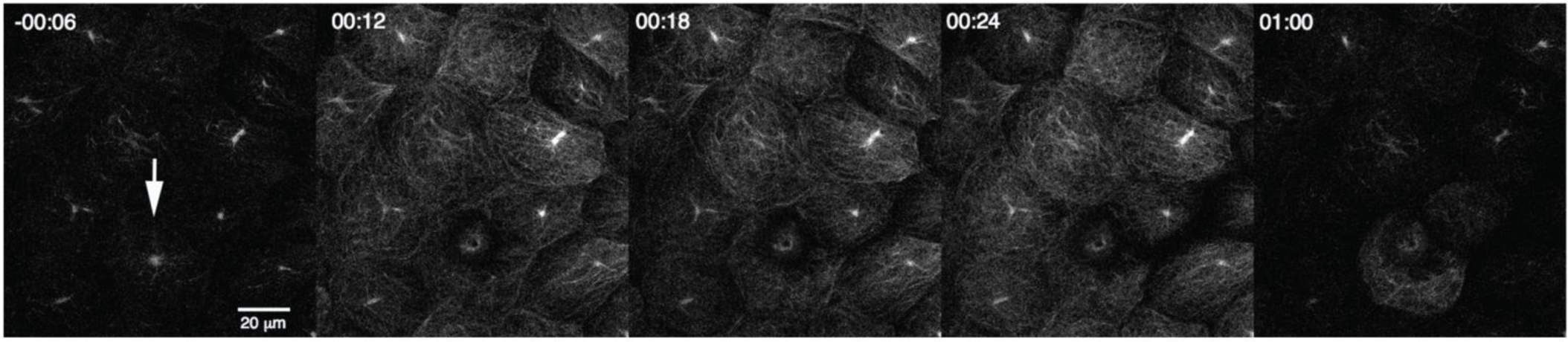
TubeCamp detects calcium increase in embryo epithelia. TubeCamp reports on microtubule-proximal changes in [Ca^2+^]_i_ in intact epithelium. Still frames from confocal movie of wounded Xenopus gastrula epidermis expressing TubeCamp. Time in min:sec; cell indicated by arrow wounded at 0 sec. Prior to wounding signal is largely confined to microtubules at presumptive base of the cilia in apical domain; wounding results in transient highlighting of entire microtubule cytoskeleton in cells neighboring the wounded cell.

**Supplementary Figure 2.**
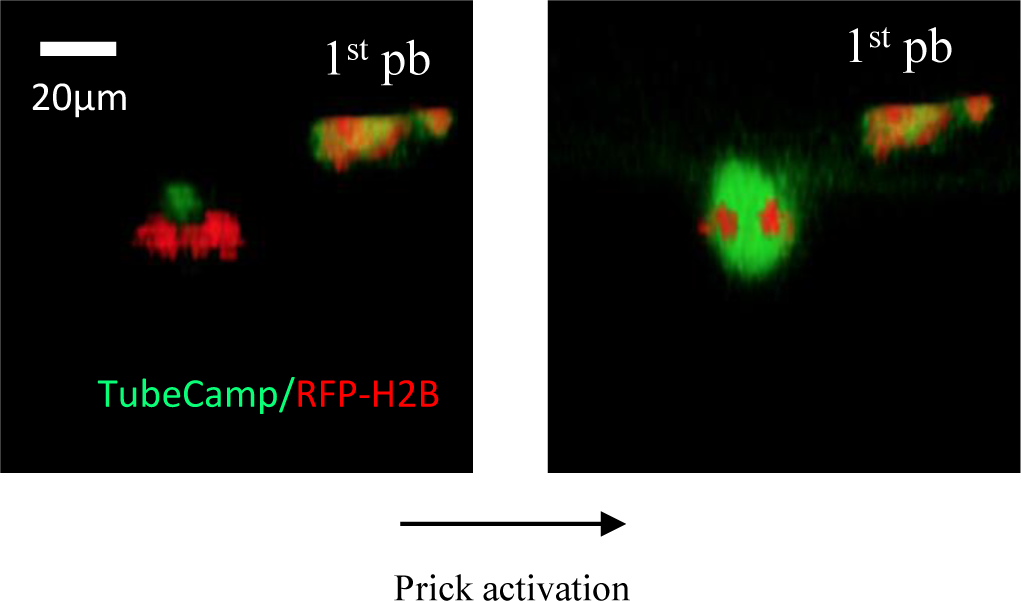
TubeCamp responds to fertilization calcium wave. Metaphase II oocytes expressing TubeCamp and RFP-H2B before (left) and 3 minutes after (right) pricking to induce fertilization-specific calcium wave. TubeCamp signal dramatically increased after pricking. 1^st^ pb: first polar body.

**Supplementary Figure 3.**
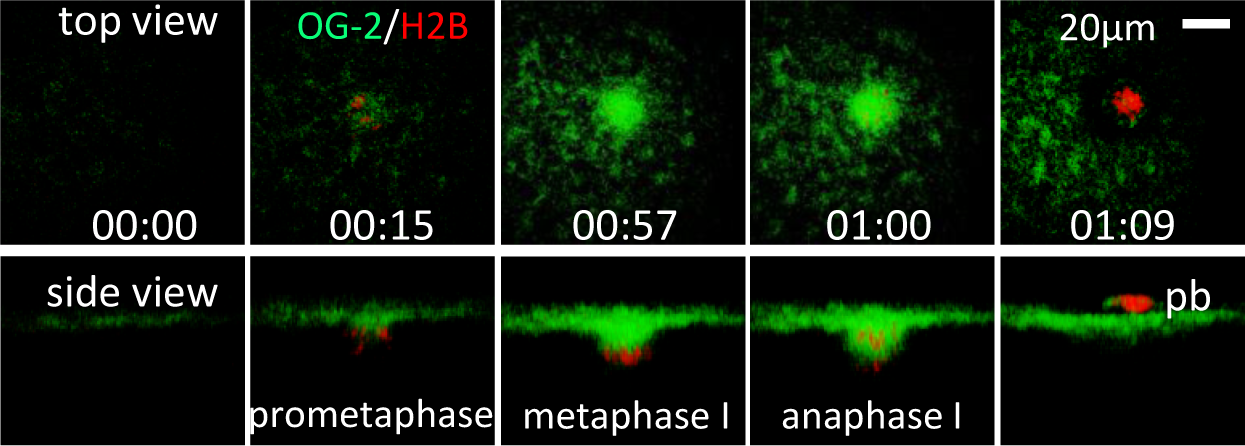
Calcium transients at the spindle assembly site. Confocal time series of an oocyte injected with Oregon-green BAPTA-2 (OG-2) and RFP-H2B at the indicated stage. Time (hr:min) is relative to the start of live cell imaging (00:00). Specific Ca^2+^ increase was seen only at the spindle assembly site, not elsewhere in the entire oocyte cortex. pb: first polar body (Movie 1).

**Supplementary Figure 4.**
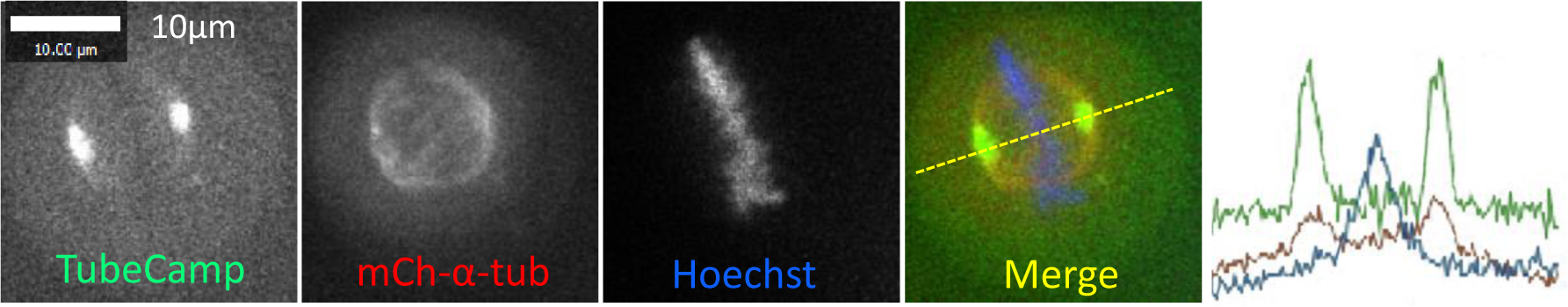
Calcium signal associated with mitotic spindles. A typical metaphase HeLa cell expressing TubeCamp and mCh-α-tubulin in the presence of Hoechst dye (blue). Left-to-right in line scan corresponds to left-to-right on the image.

**Supplementary Figure 5.**
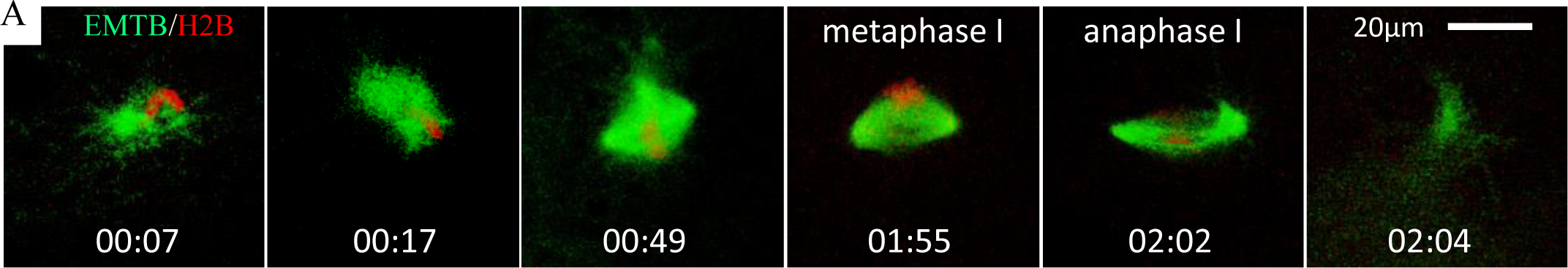
Spindle assembly in single cell aspirates. Time series of spindle formation around a single demembranated sperm in micro-aspirate derived from a single Xenopus oocyte. (Multiple droplets can be obtained from a single oocyte.) The oocyte was injected with RFP-H2B and eGFP-EMTB mRNAs and stimulated with progesterone. Cytoplasm was aspirated at the time of germinal vesicle breakdown (GVBD), mixed with demembranated sperm nuclei (00:00), and subjected to time lapse imaging. An aster formed shortly at one end of the sperm (00:07). Metaphase to anaphase transition was evident with the stretching followed by disappearance of microtubules. No chromosome segregation occurred because of the chromosomes are haploid (single sisters). Often the chromosomes moved away from the surface of the droplet after anaphase, beyond the detection limit of our confocal systems. Unlike in intact oocytes which form a metaphase II spindle after meiosis I, we have not seen any of our GVBD extracts formed a second spindle after anaphase.

## Reference List

1. Salmon, E.D. & Wolniak, S.M. Taxol stabilization of mitotic spindle microtubules: analysis using calcium-induced depolymerization. Cell Motil. 4, 155–167 (1984).

2. Pratt, M.M., Otter, T., & Salmon, E.D. Dynein-like Mg2+-ATPase in mitotic spindles isolated from sea urchin embryos (Strongylocentrotus droebachiensis). J. Cell Biol. 86, 738–745 (1980).

3. Poenie, M., Alderton, J., Steinhardt, R., & Tsien, R. Calcium rises abruptly and briefly throughout the cell at the onset of anaphase. Science 233, 886– 889 (1986).

4. Steinhardt, R.A. & Alderton, J. Intracellular free calcium rise triggers nuclear envelope breakdown in the sea urchin embryo. Nature 332, 364– 366 (1988).

5. Groigno, L. & Whitaker, M. An anaphase calcium signal controls chromosome disjunction in early sea urchin embryos. Cell 92, 193–204 (1998).

6. Sun, L. & Machaca, K. Ca(2+)(cyt) negatively regulates the initiation of oocyte maturation. J. Cell Biol. 165, 63–75 (2004).

7. Tombes, R.M. & Borisy, G.G. Intracellular free calcium and mitosis in mammalian cells: anaphase onset is calcium modulated, but is not triggered by a brief transient. J. Cell Biol. 109, 627–636 (1989).

8. Berridge, M.J. Calcium microdomains: organization and function. Cell Calcium 40, 405–412 (2006).

9. Wang, L.Y. & Augustine, G.J. Presynaptic nanodomains: a tale of two synapses. Front Cell Neurosci. 8, 455 (2014).

10. Li, R., Leblanc, J., He, K., & Liu, X.J. Spindle function in Xenopus oocytes involves possible nanodomain calcium signaling. Mol. Biol. Cell 27, 3273– 3283 (2016).

11. Tian, L. et al. Imaging neural activity in worms, flies and mice with improved GCaMP calcium indicators. Nat. Methods 6, 875–881 (2009).

12. Faire, K. et al. E-MAP-115 (ensconsin) associates dynamically with microtubules in vivo and is not a physiological modulator of microtubule dynamics. J. Cell Sci. 112 (Pt 23), 4243–4255 (1999).

13. Vasileiou, T., Foresti, D., Bayram, A., Poulikakos, D., & Ferrari, A. Toward Contactless Biology: Acoustophoretic DNA Transfection. Sci. Rep. 6, 20023 (2016).

14. von Dassow G., Verbrugghe, K.J., Miller, A.L., Sider, J.R., & Bement, W.M. Action at a distance during cytokinesis. J. Cell Biol. 187, 831–845 (2009).

15. Shao, H., Li, R., Ma, C., Chen, E., & Liu, X.J. Xenopus oocyte meiosis lacks spindle assembly checkpoint control. J. Cell Biol. 201, 191–200 (2013).

16. Zhao, Y. et al. An expanded palette of genetically encoded Ca(2)(+) indicators. Science 333, 1888–1891 (2011).

17. Mandato, C.A. & Bement, W.M. Actomyosin transports microtubules and microtubules control actomyosin recruitment during Xenopus oocyte wound healing. Curr. Biol. 13, 1096–1105 (2003).

18. Clark, A.G. et al. Integration of single and multicellular wound responses. Curr. Biol. 19, 1389–1395 (2009).

19. Busa, W.B. & Nuccitelli, R. An elevated free cytosolic Ca2+ wave follows fertilization in eggs of the frog, Xenopus laevis. J. Cell Biol. 100, 1325–1329 (1985).

20. Skoufias, D.A. et al. S-trityl-L-cysteine is a reversible, tight binding inhibitor of the human kinesin Eg5 that specifically blocks mitotic progression. J. Biol. Chem. 281, 17559–17569 (2006).

21. Dargan, S.L. & Parker, I. Buffer kinetics shape the spatiotemporal patterns of IP3-evoked Ca2+ signals. J. Physiol 553, 775–788 (2003).

22. Heald, R. et al. Self-organization of microtubules into bipolar spindles around artificial chromosomes in Xenopus egg extracts. Nature 382, 420– 425 (1996).

23. Mulligan, I.P. & Ashley, C.C. Rapid relaxation of single frog skeletal muscle fibres following laser flash photolysis of the caged calcium chelator, diazo-2. FEBS Lett. 255, 196–200 (1989).

24. Kume, S. et al. The Xenopus IP3 receptor: structure, function, and localization in oocytes and eggs. Cell 73, 555–570 (1993).

25. Ma, C. et al. Cdc42 activation couples spindle positioning to first polar body formation in oocyte maturation. Curr. Biol. 16, 214–220 (2006).

26. Turner, D.L. & Weintraub, H. Expression of achaete-scute homology 3 in Xenopus embryos converts ectodermal cells to a neural fate. Genes Dev. 8, 1434–1447 (1994).

27. Liu, X.S. & Liu, X.J. Oocyte isolation and enucleation. Methods Mol. Biol. 322, 31–41 (2006).

28. Sheng, Y. et al. A serotonin receptor antagonist induces oocyte maturation in both frogs and mice: evidence that the same G proteincoupled receptor is responsible for maintaining meiosis arrest in both species. J. Cell Physiol 202, 777–786 (2005).

29. Zhang, X. et al. Polar Body Emission Requires a RhoA Contractile Ring and Cdc42-Mediated Membrane Protrusion. Dev. Cell 15, 386–400 (2008).

30. Leblanc, J. et al. The Small GTPase Cdc42 Promotes Membrane Protrusion during Polar Body Emission via ARP2-Nucleated Actin Polymerization. Mol. Hum. Reprod. 17, 305–316 (2011).

31. Shao, H. et al. Aurora B regulates spindle bipolarity in meiosis in vertebrate oocytes. Cell Cycle 11, 2672–2680 (2012).

32. Lohka, M.J. & Masui, Y. Formation in vitro of sperm pronuclei and mitotic chromosomes induced by amphibian ooplasmic components. Science 220, 719–721 (1983).

33. Murray, A.W. Cell cycle extracts. Methods Cell Biol. 36, 581–605 (1991).

